# Ribosome display of *N*-linked glycoproteins in cell-free extracts

**DOI:** 10.1101/2022.06.09.495442

**Authors:** Sean S. Chung, Erik J. Bidstrup, Jasmine M. Hershewe, Katherine F. Warfel, Michael C. Jewett, Matthew P. DeLisa

## Abstract

Ribosome display is a powerful *in vitro* method for the selection and directed evolution of proteins expressed from combinatorial libraries. However, because ribosome display is typically performed with standard *in vitro* translation reagents, the ability to display proteins with complex post-translational modifications such as glycosylation is limited. To address this technological gap, here we developed a set of complementary methods for producing stalled ribosome complexes that displayed asparagine-linked (*N*-linked) glycoproteins in conformations amenable to downstream functional and glyco-structural interrogation. The ability to generate glycosylated ribosome-nascent chain (glycoRNC) complexes was enabled by integrating SecM-mediated translation arrest with methods for cell-free synthesis of *N*-glycoproteins. This integration yielded a novel capability for translating and displaying target proteins modified efficiently and site-specifically with different *N*-glycan structures. Moreover, the encoding mRNAs remained stably attached to stalled ribosomes both before and after biopanning, thereby providing the genotype– glycophenotype link between an arrested glycoprotein and its RNA message. We anticipate that our method will enable selection and evolution of *N*-linked glycoproteins with advantageous biological and biophysical properties.

## INTRODUCTION

Asparagine-linked (*N*-linked) protein glycosylation, the chemical modification of specific amino acid sidechains with oligosaccharides, is a conserved co- and post-translational modification (PTM) that occurs in all domains of life ^1^. As one of the most abundant PTMs ^2,3^, it serves to expand the diversity of secretory and membrane proteins ^4^ by introducing an additional information layer to these proteins. The attachment of *N*-glycans has numerous functional and structural consequences, modulating properties such as protein folding kinetics and stability, receptor binding, enzymatic activity, and localization, among others ^5,6^.

The *N*-linked glycosylation mechanism minimally involves the following: (i) assembly of *N*-glycans in the cytoplasm by sequential addition of nucleotide-activated sugars onto a lipid phosphocarrier (*e.g*., dolichylphosphate in eukaryotes and archaea or undecaprenylphosphate in bacteria ^7^); (ii) shuttling of lipid-linked oligosaccharides (LLOs) across the membrane by a flippase; and (iii) transfer of oligosaccharides from the lipid carrier onto asparagine residues in acceptor proteins by an oligosaccharyltransferase (OST) ^8–10^ in the endoplasmic reticulum (ER) of eukaryotes or in the periplasm of bacteria. A major breakthrough in our ability to study and engineer the *N*-glycosylation process occurred when Aebi and coworkers functionally transferred the bacterial glycosylation machinery encoded by the *Campylobacter jejuni pgl* locus into *Escherichia coli* cells, which do not natively perform protein glycosylation ^11^. Since this early pioneering work, diverse proteins of prokaryotic and eukaryotic origin have been *N*-glycosylated in engineered *E. coli* cells carrying the *pgl* glycosylation pathway ^12–14^ or other heterologous glycosylation pathways ^15,16^. More recently, glycoengineered *E. coli* have been leveraged as source strains to provide cell-free extracts selectively enriched with protein glycosylation machinery, including OSTs and LLOs ^17–24^. Upon addition of cofactors and plasmid DNA encoding an acceptor protein of interest, these glyco-enriched cell-free extracts enable a one-pot reaction scheme for site-specific expression and glycosylation of target glycoproteins at relatively high titers.

The creation of *E. coli* cell-based and cell-free methods for customizable protein glycosylation paves the way for development of advanced peptide/protein display techniques such as phage ^25^, ribosome ^26,27^, and mRNA ^28^ display for high-throughput screening of *N*-glycoprotein libraries. Although many methods are available for *in vitro* selection of peptides and proteins, comparatively little has been published on *in vitro* selection of glycopeptides and glycoproteins. In two notable examples, phage display was extended to include *N*-linked protein glycosylation based on the *C. jejuni pgl* system ^29,30^. In this method, genetic fusions between target *N*-glycoproteins and the minor phage coat protein g3p were expressed in glycoengineered *E. coli*, enabling production of M13 filamentous phage populations that exposed an *N*-linked glycan on their surface (glycophages). The concomitant packaging of the fusion-encoding phagemid into the glycophage particles established a physical coupling between the phenotype of the displayed glycoprotein (glycophenotype) and the corresponding genotype. The resulting genotype-glycophenotype link was subsequently leveraged to select functional glycosylation sequons from libraries of randomized acceptor sequences ^29,30^. Despite the successful history of phage display, crucial disadvantages of this methodology include: (i) the limitation of library diversity by transformation efficiency, with typical library sizes in the range of ~10^6^ to 10^9^ members; (ii) the potential for library members to be toxic to *E. coli* cells (*e.g*., becoming stalled in the Sec translocase) and thus excluded from the library; (iii) the handling of large libraries is labor intensive due to the need for infection of bacteria at several steps of the phage display cycle, including during initial production of the phage library as well as during recovery and isolation of phagemids from enriched phages; and (iv) introduction of modifications such as glycosylation can negatively impact phage infectivity.

Entirely cell-free display systems like ribosome or mRNA display have the potential to overcome these shortcomings. One benefit of these methods is that the DNA library is first transcribed *in vitro*, potentially leading to larger libraries (>10^9^) and greater diversity. In conventional ribosome display, the resulting mRNA lacks a stop codon, giving rise upon translation to stable mRNA–ribosome–protein complexes, which can be directly used for selection against an immobilized target. The resulting mRNA is obtained upon dissociating the ribosomal subunits, reverse transcribed and amplified for the next round. In mRNA display, the transcribed mRNA is first ligated to a DNA linker connected to puromycin, leading to stalling of the ribosome at the RNA–DNA junction when the mRNA is translated *in vitro*.

While these methods have been widely used with great success, to date there have been no reports of extending the scope of ribosome or mRNA display to include *N*-linked glycoproteins. This technology gap is most likely because conventional *in vitro* translation systems, which these methods depend on for producing displayed proteins, are limited by their inability to co-activate efficient protein synthesis and glycosylation. For example, the best characterized and most widely used cell-free protein synthesis (CFPS) systems based on *E. coli* S30 extract are incapable of making glycoproteins because the *E. coli* source strains lack endogenous glycosylation machinery ^31^. Likewise, the PURE system, a completely reconstituted cell-free system composed of 36 *E. coli* enzymes that are involved in transcription and translation as well as highly purified 70S ribosomes ^32^, is devoid of glycosylation machinery. Other commonly used eukaryotic CFPS systems including those based on rabbit reticulocyte and wheat germ extracts also cannot perform glycosylation because they lack microsomes ^33^. While glycosylation can be introduced via supplementation with microsomes (*e.g*., canine pancreas microsomes), the resulting glycoprotein yields in these systems are often low due to the poor compatibility between the extract translational machinery and microsomal glycosylation machinery ^34,35^. Moreover, microsome-mediated glycosylation requires translocation of target proteins into microsomal vesicles which is likely incompatible with protein stalling on ribosomes due to the sequestration of glycoproteins inside the vesicles. Indeed, the glycophage display method described above was only possible because the processes of phage assembly and *N*-linked protein glycosylation could be harmonized in the periplasm of living *E. coli* cells.

In this study, we investigated the extent to which ribosome display was compatible with emerging methods for cell-free glycoprotein biosynthesis (CFGpS) based on extracts derived from glycoengineered *E. coli* ^17,18^, which effectively couple transcription/translation with glycosylation. Like other screening methods, a prerequisite for the selection of proteins from ribosome display libraries is the genotype-phenotype link, which is accomplished during *in vitro* translation by stabilizing a complex consisting of the ribosome, the mRNA, and the nascent, correctly folded polypeptide. Here, we hypothesized that stalled proteins could be glycosylated by *C. jejuni*. glycosylation machinery, and furthermore that CFGpS could be functionally integrated with the translation and stalling processes. To test this hypothesis, we attempted to create a genotype-glycophenotype link using the SecM-mediated translation arrest mechanism to stall glycoproteins and their encoding mRNA on ribosomes ^36^. Previous studies demonstrated that when the 17-amino acid stall sequence (FSTPVWISQAQGIRAGP) derived from *E. coli* SecM (SecM17) was expressed at the C-terminus of a protein of interest (POI), it promoted stalling of ribosomes that were stably tethered to the heterologous POI and its encoding mRNA both in intact *E. coli* cells and their cell-free extracts ^37,38^. By combining this same SecM17 stall sequence with cell-free glycosylation strategies, we produced glycosylated ribosome-nascent chain (glycoRNC) complexes that displayed several different *N*-linked glycoproteins in conformations that were sufficiently exposed to permit downstream functional and glyco-structural interrogation. Importantly, mRNA transcripts encoding the displayed glycoproteins were found stably attached to stalled ribosomes both before and after biopanning, thereby providing the crucial physical linkage between an arrested glycoprotein and its RNA transcript, paving the way for future selection and evolution of *N*-linked glycoproteins with desirable properties.

## RESULTS

### SecMI7-mediated display of *N*-linked glycoproteins on stalled ribosomes

The engineering of ribosome display for *N*-linked glycoproteins hinges on generation of glycoRNC complexes (**Fig. 1**). To this end, we first investigated whether target acceptor proteins were amenable to glycosylation post-stalling using a hybrid *in vivo*-*in vitro* approach. For the *in vivo* step, expression of a POI-SecM17 fusion was performed in living *E. coli* cells from which 70S ribosomes were subsequently isolated. For the *in vitro* step, isolated 70S ribosomes were subjected to an *in vitro* reconstituted glycosylation system to install *N*-glycans on the SecM17-arrested POI. A pET28a-based expression plasmid was constructed that enabled fusions between any POI and the SecM17 stall sequence (**Fig. 1**). Importantly, we used a previously identified linker sequence between the POI and SecM17 that had sufficient length to fully expose stalled proteins outside the ribosome exit tunnel, rendering them available for functional interrogation ^37^. Our initial POI was the *E. coli* immunity protein, Im7, a globular 87-residue protein that is well-expressed in *E. coli*. While not a native glycoprotein, Im7 has been shown to tolerate *N*-glycan installation at numerous artificial DQNAT acceptor sites throughout its structure using both cellular and cell-free glycosylation systems based on the *C. jejuni pgl* machinery ^14^. We specifically chose to focus on the Im7^N58^ mutant (where the superscript denotes the location of the asparagine residue) as it is efficiently glycosylated *in vitro*^14^.

**Figure 1.**
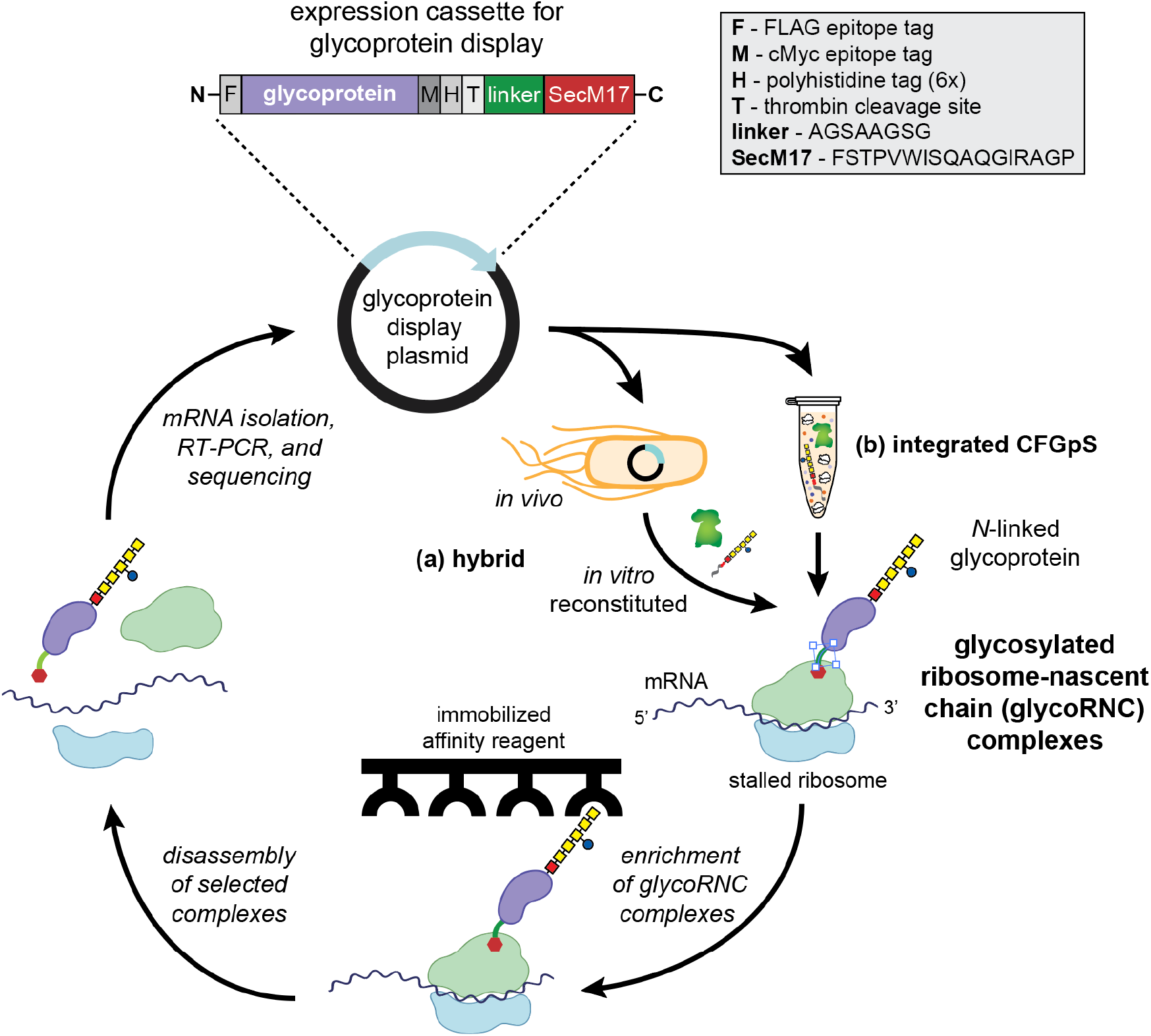
Cell-free glycoprotein display on ribosomes. Schematic showing the principle of ribosome display for *N*-linked glycoproteins, a protein engineering method that enables selection and directed evolution of functional and structural structure-function properties of glycoproteins. (top) First, the DNA encoding a glycoprotein library is introduced into plasmid pET28a (for *in vivo* expression) or pJL1 (for *in vitro* expression) in which a flexible linker sequence (e.g., AGSAAGSG) and SecM17 stall sequence (FSTPVWISQAQGIRAGP) are introduced at the C-terminus, along with additional epitope tags for detection and purification. From here, there are two possible routes for making ribosome-stalled glycoprotein libraries. The first involves transformation of the plasmid DNA library into *E. coli* cells and expression of glycoprotein–SecM17 fusions, followed by isolation of 70S ribosomes from cells. In this hybrid approach, glycoprotein mRNA is transcribed and translated entirely *in vivo*, while installation of *N*-glycans on ribosome-stalled glycoproteins is performed in a subsequent *in vitro* glycosylation step. The second route involves an entirely cell-free process in which the plasmid DNA library is used to prime an integrated cell-free glycoprotein synthesis (CFGpS) reaction mixture in which transcription and translation are coupled to *N*-linked glycosylation in a single pot. Both routes yield glycosylated ribosome-nascent chain (glycoRNC) complexes in which glycoproteins are tethered to 70S ribosomes along with the encoding mRNA (right side). The desired glycoRNC complexes are then subjected to function-based affinity selection (*e.g*., binding to immobilized antigen) and/or glycosylation-based affinity selection (*e.g*., binding to immobilized antibody or lectin that specifically recognizes the *N*-glycan). Non-specific glycoRNC complexes are removed by intensive washing and bound complexes are dissociated by EDTA (or specifically eluted with antigen or glycan). RNA is isolated from the dissociated complexes and reverse transcribed to cDNA. The resulting cDNA is amplified by PCR and the PCR product is then used for the next cycle of enrichment (a portion can be analyzed by cloning and sequencing and/or by ELISA). Image created with biorender.com.

To confirm ribosome display of Im7^N58^–SecM17 fusions, we first isolated 70S ribosomes from *E. coli* cell lysates via sucrose cushion centrifugation. Consistent with earlier findings that SecM17 mediates the display of heterologous proteins on intact ribosomes ^37,38^, Western blot analysis of the resulting sucrose gradient fractions demonstrated that only the Im7^N58^–SecM17 fusion but not Im7^N58^ lacking the SecM17 stall sequence was present in ribosome fractions, as judged by the absorbance profile at 254 nm (A254) and sedimentation position (70S) (**Supplementary Fig. 1a**). These results confirmed that co-elution of Im7^N58^ with ribosomes depended on the presence of the SecM stall sequence. Expression and arrest of Im7^N58^–SecM17 on ribosomes did not significantly affect ribosomal composition, as evidenced by the nearly identical protein profiles for ribosome preparations derived from *E. coli* BL21(DE3) cells expressing Im7^N58^ with or without the SecM17 stall sequence and BL21(DE3) cells carrying an empty expression vector (**Supplementary Fig. 1b**). Likewise, the growth rates of cells expressing the Im7^N58^–SecM17 fusion and unfused Im7^N58^ were indistinguishable from the growth rate of cells containing an empty expression vector during the 30–90 min induction period (data not shown).

### Generation of glycoRNC complexes by *in vitro* reconstituted glycosylation system

To create glycoRNC complexes, we hypothesized that 70S ribosome preparations derived from the *in vivo* step above could be used as purified acceptor substrates for an *in vitro* reconstituted glycosylation assay. To test this notion, RNC complexes displaying aglycosylated Im7^N58^–SecM17 were combined with purified OST enzyme, namely PglB from *C. jejuni* (*Cj*PglB), and solvent-extracted lipid-linked oligosaccharides bearing the *C. jejuni M*-glycan (*Cj*LLOs) (**Fig. 2a**). Western blotting was performed on the *in vitro* glycosylation reaction products using an anti-FLAG antibody to detect the Im7^N58^–SecM17 protein and hR6 serum to specifically detect the *C. jejuni N*-glycan ^39^. A shift in the apparent molecular weight of Im7^N58^–SecM17 in the anti-FLAG immunoblot that corresponded to a similarly sized band detected in the hR6 immunoblot confirmed that ribosome-tethered Im7^N58^–SecM17 was nearly 100% glycosylated under the conditions tested (**Fig. 2b**). When *Cj*PglB was omitted from the reaction, we observed no detectable glycosylation of ribosome-stalled Im7^N58^–SecM17, confirming the expected OST-dependent installation of *N*-glycans onto ribosome-stalled acceptor proteins.

**Figure 2.**
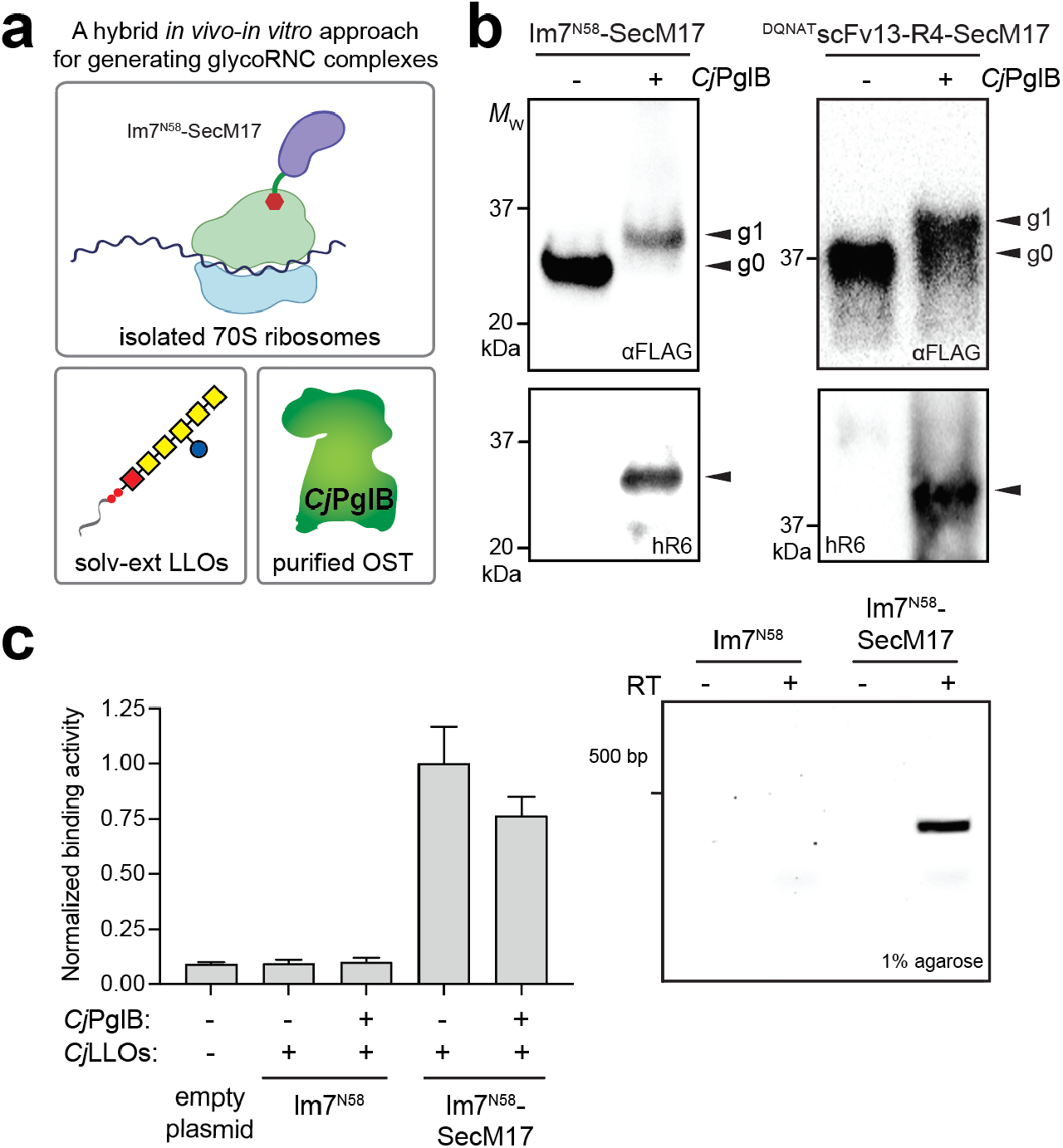
A hybrid *in vivo-in vitro* strategy for generating and selecting glycoRNC complexes. (a) Schematic of hybrid strategy in which 70S ribosomes displaying aglycosylated target proteins are produced using living cells (*in vivo*) and, following isolation from cells, subjected to cell-free glycosylation involving purified *Cj*PglB and solvent-extracted *Cj*LLOs (*in vitro*). Image created with biorender.com (b) Western blot analysis of 70S ribosome fractions derived from cells expressing Im7^N58^–SecM17 or ^DQNAT^scFv13-R4– SecM17 and subjected to *in vitro* glycosylation in the presence (+) or absence (−) of *Cj*PglB. Blots were probed with anti-FLAG antibody to detect acceptor protein (top) and hR6 serum against the glycan (bottom). Markers for aglycosylated (g0) and singly glycosylated (g1) forms of acceptor proteins are indicated on right. Molecular weight (*M_W_*) markers are indicated on left. Western blots are representative of biological replicates (*n* = 2). (c) ColE7-binding activity measured for 70S ribosomes isolated from cells expressing Im7^N58^ or Im7^N58^–SecM17 in the presence (+) or absence (−) of *Cj*PglB. 70S ribosomes from cells carrying empty plasmid served as the negative control. Data are the average of biological replicates (*n* = 3) and error bars represent the standard deviation. (d) Post-selection detection of mRNA associated with ribosomes enriched via ColE7 biopanning. mRNA isolated from functionally selected ribosomes corresponding to glycosylated Im7^N58^ and Im7^N58^–SecM17 was subjected to RT-PCR in the presence (+) or absence (−) of reverse transcriptase (RT). Gel results are representative of biological replicates (*n* = 2).

To demonstrate the modularity of our hybrid approach, we swapped the POI in the pET28a-based expression plasmid from Im7^N58^ to a single-chain Fv antibody specific for β-galactosidase modified with a DQNAT glycosylation tag at its N-terminus (^DQNAT^scFv13-R4) ^40^. It should be noted that scFv13-R4 bearing a C-terminal DQNAT tag was previously found to be amenable to glycosylation by the *C. jejuni* glycosylation machinery ^12,17^. Here, the ^DQNAT^scFv13-R4-SecM17 construct was present in sucrose gradient fractions containing 70S ribosomes, as judged by the absorbance profile at 254 nm (A254), the sedimentation position, and Western blot analysis of the resulting sucrose gradient fractions (**Supplementary Fig. 2**). Similar to Im7^N58^–SecM17, ^DQNAT^scFv13-R4-SecM17 was efficiently glycosylated in a *Cj*PglB-dependent manner (**Fig. 2b**), leading to the formation of glycoRNC complexes displaying the glycosylated scFv antibody. We next determined whether intact mRNA encoding the POI remained associated with glycoRNC complexes following functional selection. To this end, a bio-panning procedure was performed in whcih *in vitro* glycosylation reaction products were directly incubated in enzyme-linked immunosorbent assay (ELISA) plates coated with ColE7, a 60-kDa bacterial toxin that is inhibited by Im7 binding ^41^. Following extensive washing, strong antigen-binding activity was measured for stalled RNC complexes displaying glycosylated Im7^N58^–SecM17 that was comparable to that observed for aglycosylated Im7^N58^–SecM17 (**Fig. 2c**), indicating that our procedures for stalling and glycosylating Im7^N58^ on ribosomes were compatible with its ColE7-binding activity. Upon verifying affinity capture, the bound glycoRNC complexes were dissociated with EDTA and the ribosome-associated RNA, including stalled mRNA, was isolated from dissociated complexes. To detect the presence of mRNA encoding Im7^N58^, reverse transcription polymerase chain reaction (RT-PCR) was performed using primers specific for the Im7^N58^ gene sequence. By this method, we determined that RNA recovered from functionally selected glycoRNC complexes gave rise to a clearly detectable amplicon corresponding in size to the expected full-length Im7^N58^ sequence (**Fig. 2d**), which was confirmed by sequencing to encode Im7^N58^. In contrast, RT-PCR performed on RNA recovered from samples corresponding to Im7^58^ lacking the SecM17 stall sequence produced no detectable amplicons, indicating that the SecM17 stall sequence is essential for functional selection of glycoRNC complexes. Collectively, these data confirmed that the model glycoprotein used in these experiments, Im7^58^, and its encoding mRNA remained stably attached to stalled ribosomes, thereby creating the essential genotype to phenotype link that is required for library screening.

### Generation of glycoRNC complexes by a one-pot cell-free method

Alongside the hybrid approach described above, we also investigated an alternative strategy for generating glycoRNC complexes via an entirely cell-free procedure that circumvented the need for *in vivo* expression and purification of stalled 70S ribosome-POI complexes as well as the supplementation of laboriously extracted/purified glycosylation components. We hypothesized that such an approach would be possible by integrating SecM-mediated translation arrest with a previously described cell-free glycoprotein synthesis (CFGpS) technology in which transcription and translation are coupled to *N*-linked glycosylation in a single pot ^17^ (Fig. 3a). To test this notion, we once again focused on the Im7^N58^ variant as a model POI and generated a cell-free expression plasmid in which this acceptor-SecM17 protein was fused to the SecM17 stall sequence in plasmid pJL1 ^42^. In parallel, S12 extracts were prepared from *E. coli* CLM24 cells that simultaneously overexpressed *Cj*PglB and the *C. jejuni* glycan biosynthesis enzymes. Our previous work demonstrated that such extracts are selectively enriched with both *Cj*PglB and *Cj*LLOS^17^ and that S12 extract can achieve higher glycosylation efficiencies than S30 extract due to its increased concentration of vesicle-bound glycosylation machinery ^19^. To generate glycoRNC complexes, batch-mode sequential CFGpS reactions were performed by priming glycoenriched extracts with plasmid DNA encoding the Im7^N58^–SecM17 construct, which induced translation and subsequent stalling of Im7^N58^–SecM17 on 70S ribosomes. Next, 70S ribosomes were isolated from CFGpS reaction mixtures by sucrose cushion centrifugation and subjected to Western blot analysis with anti-FLAG and hR6 serum antibodies. As was seen above for *in vitro* reconstituted glycosylation, the CFGpS reaction mixture promoted efficient glycosylation of Im7^N58^-SecM17 on stalled ribosomes, with nearly 100% of the acceptor protein appearing as the g1 form in the anti-FLAG blot (**Fig. 3b**). As expected, a control experiment performed with S12 extracts enriched with only *CjLLOS* but not *Cj*PglB resulted in no detectable glycosylation of Im7^N58^–SecM17 on stalled ribosomes. Importantly, the mRNA encoding SecM17-stalled Im7^N58^ was stably attached to ribosomes as evidenced by the prominent amplicon generated via RT-PCR of ribosome-associated RNA that was recovered from RNC complexes displaying glycosylated Im7^N58^–SecM17 (**Fig. 3c**). Nearly identical results were obtained when CFGpS reactions were primed with plasmid DNA encoding a different acceptor POI, namely an scFv antibody specific for human epidermal growth factor receptor 2 (scFv-HER2) that was modified at its C-terminus with a DQNAT glycosylation tag ^14,43^ (**Fig. 3b and c**). Using immobilized HER2 as antigen, we selected glycoRNC complexes displaying glycosylated scFv-HER2^DQNAT^–SecM17 and observed that the encoding mRNA remained associated with these functionally selected glycoRNCs (**Fig. 3d**). In contrast, no mRNA was detected for ribosome complexes prepared from CFGpS reactions primed with plasmid DNA encoding unfused scFv-HER2^DQNAT^, indicating that functional selection of glycoRNCs was dependent on the SecM17 stall sequence.

**Figure 3.**
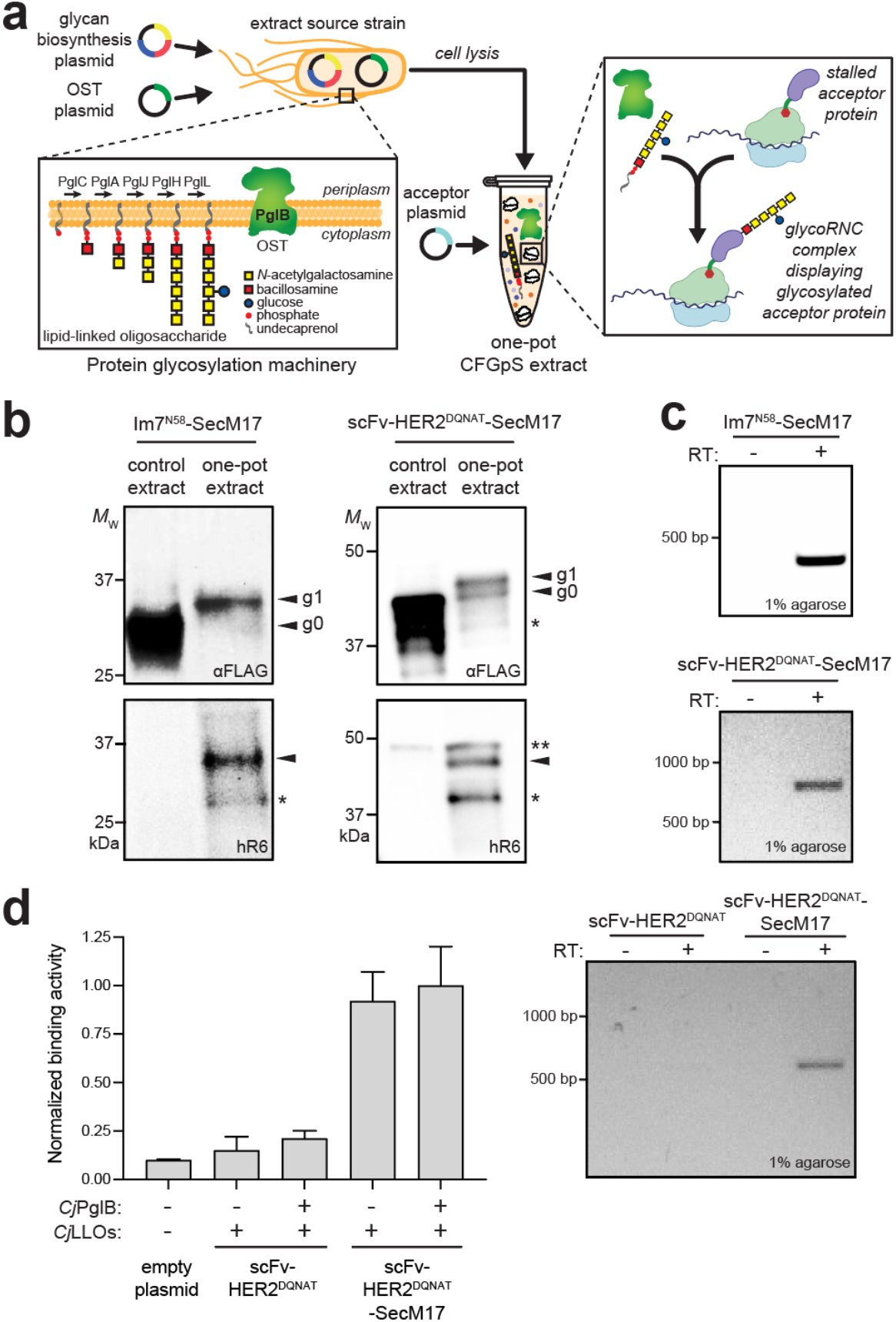
A one-pot cell-free strategy for generating and selecting glycoRNC complexes. (a) Schematic of one-pot strategy in which 70S ribosomes displaying glycosylated target proteins are produced using CFGpS involving extracts co-enriched with *Cj*PglB and *Cj*LLOs. Image created with biorender.com (b) Western blot analysis of 70S ribosome fractions derived from CFGpS extracts co-enriched with *Cj*PglB and *Cj*LLOs (one-pot) or enriched with only *Cj*LLOs (control) that were primed with plasmids encoding Im7^N58^–SecM17 or scFv-HER2^DQNAT^–SecM17. Blots were probed with anti-FLAG antibody to detect acceptor protein (top) and hR6 serum against the glycan (bottom). Markers for aglycosylated (g0) and singly glycosylated (g1) forms of acceptor proteins are indicated on right. Asterisk (*) denotes degradation product; two asterisks (**) denotes non-specific product. Molecular weight (*M_W_*) markers are indicated at left. Western blots are representative of biological replicates (*n* = 2). (c) Pre-selection detection of mRNA associated with 70S ribosomes displaying glycosylated Im7^N58^–SecM17 or scFv-HER2^DQNAT^–SecM17. RT-PCR was performed in the presence (+) or absence (−) of RT. (d, left) HER2-binding activity measured for 70S ribosomes isolated from CFGpS reactions primed with plasmid DNA encoding scFv-HER2^DQNAT^ or scFv-HER2^DQNAT^–SecM17 in the presence (+) or absence (−) of *Cj*PglB. 70S ribosomes from cells carrying empty plasmid served as the negative control. Data are the average of biological replicates (*n* = 3) and error bars represent the standard deviation. (d, right) Post-selection detection of mRNA associated with ribosomes enriched via HER2 biopanning. mRNA isolated from functionally selected ribosome complexes corresponding to glycosylated scFv-HER2^DQNAT^ or scFv-HER2^DQNAT^–SecM17 was subjected to RT-PCR in the presence (+) or absence (−) of RT. Gel results are representative of biological replicates (*n* = 2).

### SecM-mediated display and glycan-mediated enrichment of a bioconjugate vaccine

To further demonstrate the utility of our glycoprotein display method, we designed a one-pot cell-free bioconjugation strategy for displaying conjugate vaccine candidates on 70S ribosomes (**Fig. 4a**). Cell-free bioconjugation is a recently described approach that leverages glyco-competent cell-free extracts for producing conjugate vaccines that retain native immunogenic structures ^20^. In this approach, cell-free extracts are co-enriched with an OST (*e.g*., *Cj*PglB) and LLOs bearing heterologously expressed capsular polysaccharides (CPS) or O-antigen polysaccharides (O-PS). The resulting extracts are then primed with plasmid DNA encoding a vaccine carrier protein such as non-acylated *Haemophilus influenzae* protein D (PD), yielding product proteins that are site-specifically glycosylated with CPS or O-PS antigens of interest.

**Figure 4.**
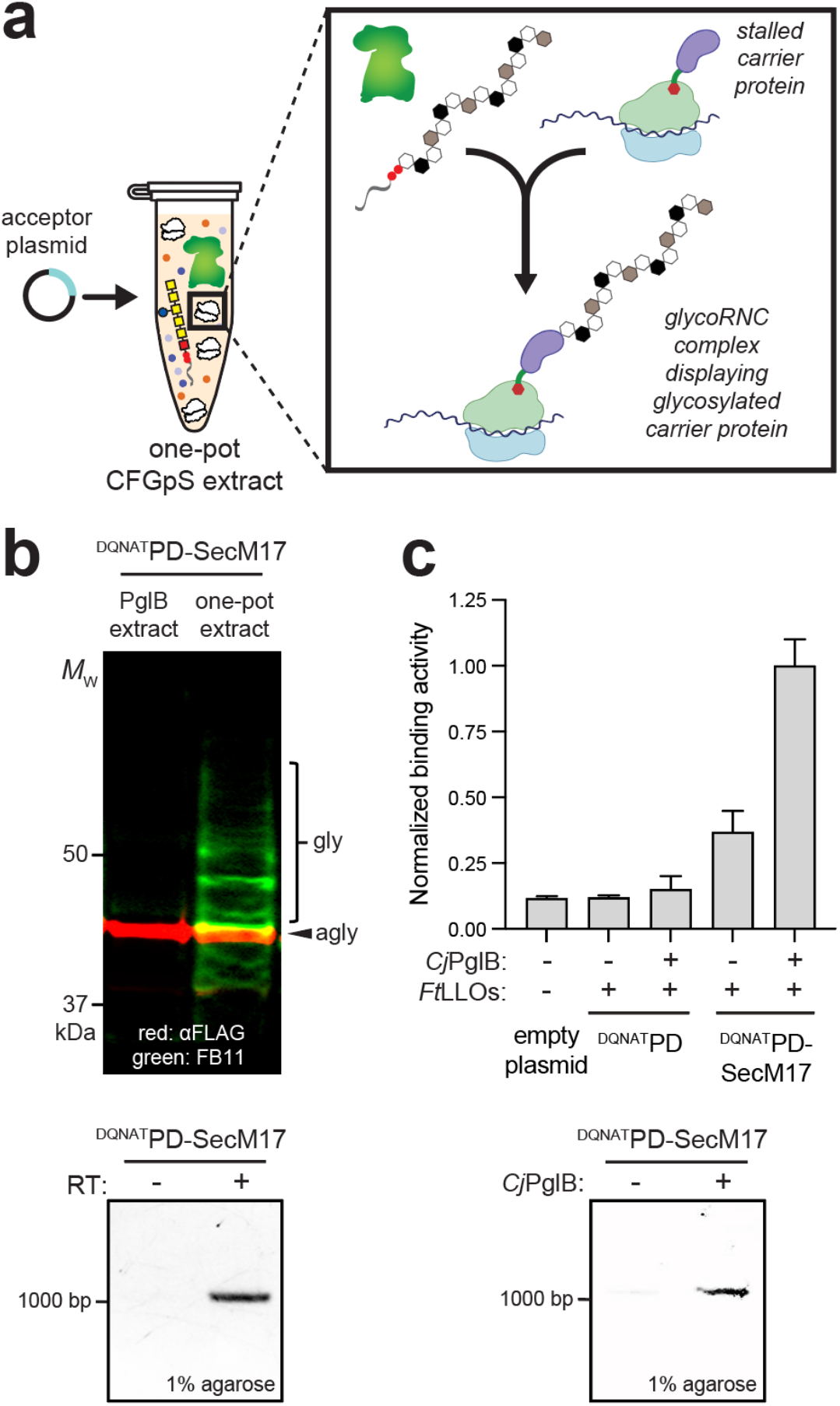
Display and selection of ribosome-stalled bioconjugate vaccine. (a) Schematic of one-pot strategy in which 70S ribosomes displaying glycosylated vaccine carrier protein (*e.g*., protein D (PD)) are produced using CFGpS involving extracts co-enriched with *Cj*PglB and *Ft*LLOs. Image created with biorender.com (b, top) Western blot analysis of 70S ribosome fractions derived from CFGpS extracts coenriched with *Cj*PglB and *Ft*LLOs (one-pot) or enriched with only *Ft*LLOs (control) that were primed with plasmid DNA encoding ^DQNAT^PD–SecM17. Merged image of blots that were probed with anti-FLAG antibody to detect the acceptor protein (red) and hR6 serum against the glycan (green). Markers for aglycosylated (agly) and multiply glycosylated (gly) forms of acceptor proteins are indicated on right. Western blots are representative of biological replicates (*n* = 2).

Here, we integrated this cell-free vaccine expression technology with SecM-mediated stalling to engineer glycoRNC complexes that displayed a vaccine carrier protein modified with an O-PS antigen. We focused on PD because it has been shown to be a safe and effective conjugate vaccine carrier protein ^44,45^ that can be efficiently glycosylated with the *Francisella tularensis* O-PS (*Ft*O-PS) antigen in CFGpS ^20^. Here, PD was modified at its N-terminus with a DQNAT acceptor motif and at its C-terminus with the SecM17 stall sequence in plasmid pJL1. In parallel, S12 extracts were prepared from *E. coli* CLM24 cells that heterologously expressed *Cj*PglB and the *Ft*O-PS biosynthesis enzymes. We have previously found that such extracts are selectively enriched with both *Cj*PglB and LLOs bearing the *Ft*O-PS antigen (*Ft*LLOs) ^20^. Next, batch-mode sequential CFGpS reactions were performed by priming these glyco-enriched extracts with plasmid DNA encoding the ^DQNAT^PD–SecM17 construct. Isolation of 70S ribosomes and Western blot analysis were performed as described above but with antibody FB11, which specifically recognizes the *Ft*O-PS antigen ^46^, in place of hR6 serum. Importantly, the CFGpS reaction mixture promoted efficient glycosylation of ^DQNAT^PD-SecM17 on stalled ribosomes with a ladder-like banding pattern (**Fig. 4b**), which was characteristic of the laddering seen in prior *Ft*O-PS bioconjugates and arises due to O-PS chain length variability through the action of the Wzy polymerase ^20,47,48^. As expected, no evidence for glycosylation was observed for extracts enriched with only *Ft*LLOs but lacking *Cj*PglB. Once again, the encoding mRNA was stably associated with glycoRNC complexes displaying stalled ^DQNAT^PD-SecM17 as evidenced by the amplicon generated via RT-PCR of ribosome-associated RNA (**Fig. 4b**). Finally, we demonstrated (b, bottom) Pre-selection detection of mRNA associated with 70S ribosomes displaying glycosylated ^DQNAT^PD–SecM17 in the presence (+) or absence (−) of RT. (c, top) FB11-binding activity measured for 70S ribosomes isolated from CFGpS reactions primed with plasmid DNA encoding ^DQNAT^PD–SecM17 in the presence (+) or absence (−) of *Cj*PglB. 70S ribosomes derived from cells carrying empty plasmid served as negative control. Data are the average of biological replicates (*n* = 3) and error bars represent the standard deviation. (c. bottom) Post-selection detection of mRNA associated with ribosomes enriched via *FtO*-PS-specific biopanning with FB11 antibody. Ribosome-associated mRNA was isolated from selected ribosome complexes corresponding to glycosylated (+) or aglycosylated (−) ^DQNAT^PD–SecM17 and subjected to RT-PCR. Gel results are representative of biological replicates (*n* = 2). selection of RNC complexes displaying glycosylated ^DQNAT^PD-SecM17 using immobilized FB11 antibody, which preferentially enriched for RNC complexes bearing the *Ft*O-PS antigen over those that were aglycosylated or those corresponding to ^DQNAT^PD that lacked the SecM17 stall sequence (**Fig. 4c**). Importantly, the encoding mRNA remained associated with the enriched glycoRNCs (**Fig. 4d**), indicating that ^DQNAT^PD-SecM17-stalled ribosomes could be directly selected based on their glyco-phenotype. In contrast, only a faint mRNA signal was detected for ribosomes prepared from CFGpS extracts that lacked *Cj*PglB, confirming that *Ft*O-PS-based enrichment had occurred.

## DISCUSSION

In this study, we showed that ribosome display was compatible with emerging methods for cell-free glycoprotein biosynthesis (CFGpS) ^17,18^, leading to the first demonstration of ribosome display of *N*-linked glycoproteins. This was accomplished by combining the SecM17 stall sequence with two complementary cell-free glycosylation strategies based on the *C. jejuni N*-glycosylation pathway. The result was the formation of stable glycoRNC complexes that displayed several different *N*-linked glycoproteins in conformations that were compatible with downstream functional and glyco-structural interrogation. While this was the first demonstration of ribosome display of *N*-linked glycoproteins, we note that the scope of phage display has previously been extended to the display of posttranslational modifications, including *N*-linked glycoproteins ^29,30^ as well as phosphorylation and phosphopantetheinylation ^49^. It should also be noted that a handful of reports have described the use of phage or mRNA to display short peptides (5-33 amino acids) that were chemically modified by either a single mannose monosaccharide or a high-mannose glycan ^50–52^. In each case, the monosaccharide or glycan attachment was through a variety of unnatural linkages including: formation of a disulfide bond between the side chain of a cysteine residue and 2-(3-nitropyridyl disulfide ethyl)-mannopyranoside (Man-Npys) ^50^; oxime ligation following oxidation of an N-terminal Ser/Thr residue ^51^; and installation of the unnatural amino acid homopropargylglycine that was “click”-glycosylated with Man9-azide through copper-assisted azide alkyne cycloaddition (CuAAAC) ^52^. In contrast, our method involved installation of *N*-linked glycans via OST-mediated formation of glycosidic bonds between the reducing-end sugar and the sidechain of an asparagine residue.

An important facet of this work was the observation that a successful genotype-glycophenotype link was created upon integration of cell-free *N*-linked protein glycosylation with stable ribosome stalling via SecM translation arrest. While not directly demonstrated here, this linkage will be crucial for future efforts focused on uncovering the functional and structural consequences of *N*-glycan installation as well as the selection and evolution of *N*-linked glycoproteins with desirable properties. For example, ribosomal stalling of glycoproteins both *in vivo* and *in vitro* may allow control of the structural and temporal context of *N*-glycan installation and subsequent protein folding, which could shed light on long-standing questions about the role of *N*-linked glycans in protein folding *in vivo*^53,54^ and the mechanism of glycosylation in CFGPS ^34^. Importantly, ribosome display of *N*-linked glycoproteins stands alongside a growing list of screening and selection tools that are amenable to glycoprotein engineering. Besides the glycophage display methods discussed above, other high-throughput genetic assays for *N*-linked glycosylation that have been described include enzyme-linked immunosorbent assay (ELISA)-based detection of periplasmic *N*-glycoproteins ^55,56^, cell-surface display of *N*-glycans and *N*-glycoproteins ^13,16,57^, and a colony replica blotting strategy called glycoSNAP (<u>glyco</u>sylation of <u>s</u>ecreted <u>*N*</u>-linked <u>a</u>cceptor <u>p</u>roteins) ^12,14,58^. Collectively, these assays are enabling the creation and evaluation of an unprecedentedly large number of intact glycoproteins (>150 in one particular study alone ^14^), for which the structure-activity relationships associated with *N*-glycan installation can be systematically catalogued or technologically exploited. Indeed, our demonstration that a bioconjugate vaccine candidate comprised of an FDA-approved carrier protein and a pathogen-specific polysaccharide was compatible with ribosome display paves the way for studying and engineering the biological, biophysical, and immunological properties of this important class of man-made glycoproteins.

## MATERIALS AND METHODS

### Bacterial strains and plasmids

*E. coli* strain DH5α was used for cloning and maintenance of plasmids. *E. coli* strain BL21(DE3) was used for *in vivo* production of RNC complexes displaying different POIs and for production of ColE7^H569A^ as described ^14^. *E. coli* strain CLM24 was used for purification of the *Cj*PglB enzyme and organic solvent-based extraction of *CjLLOs* used for *in vitro* glycosylation as described ^17^. CLM24 was also used as the source strain for preparing cell-free extracts with selectively enriched glycosylation components as described ^17^. CLM24 is a glyco-optimized derivative of W3110 that carries a deletion in the gene encoding the WaaL ligase, thus facilitating the accumulation of preassembled glycans on undecaprenylphosphate ^15^.

Plasmids constructed in this study were made using standard cloning protocols and included the following. For *in vivo* expression, plasmid pET–Im7^N58^–SecM17 was constructed by inserting the PCR-amplified product encoding Im7^N58^ in place of the gene encoding scFv13–R4 in plasmid pET–scFv13–R4–SecM17 ^37^. Plasmid pET–^DQNAT^scFv13–R4–SecM17 was constructed by similar replacement but with a PCR-amplified product encoding ^DQNAT^scFv13-R4 in which the N-terminal DQNAT motif was introduced via the forward primer. Plasmid pET–Im7^N58^ was constructed by PCR amplification of the gene encoding Im7^N58^ from plasmid pTrc99S–YebF–Im7^N58 14^ and subsequent Gibson isothermal assembly ^59^ of the resulting PCR product into the amplified plasmid backbone of pET28a. For *in vitro* expression, plasmid pJL1–Im7^N58^–SecM17 was constructed by PCR amplification of the gene encoding Im7^N58^–SecM17 from plasmid pET–Im7^N58^–SecM17 and Gibson assembly of the resulting PCR product into the amplified pJL1 plasmid backbone. Plasmids pJL1–scFv–HER2^DQNAT^–SecM17 and pJL1–^DQNAT^PD–SecM17 were constructed by PCR amplification of the genes encoding scFv– HER2^DQNAT^ from plasmid pTrc99S–YebF–scFv–HER2^DQNAT^ ^14^ and PD from plasmid pJL1–PD^4xDQNAT^ ^20^, respectively, and inserting the resulting PCR products in place of Im7^N58^ in amplified plasmid pJL1–Im7^N58^–SecM17 via Gibson assembly. The same PCR products were used to construct plasmids pJL1–scFv–HER2^DQNAT^ and pJL1–^DQNAT^PD by Gibson assembly into plasmid pJL1. Additional plasmids constructed previously and used in this study included: plasmid pSN18 for expression and purification of *Cj*PglB ^60^; plasmid pSF–*Cj*PglB for selective enrichment of *Cj*PglB in cell-free extracts ^61^; plasmid pMW07–pglΔB for expression and extraction of *Cj*LLOs ^12^; and plasmid pET28– ColE7^H569A^ for expression and purification of ColE7 ^14^. All plasmids were confirmed by DNA sequencing at the Biotechnology Resource Center (BRC) Genomics Facility (RRID:SCR_021727) in the Cornell Institute of Biotechnology.

### Ribosome isolation

Cells transformed with the pET28a-derived constructs were grown in 100-mL cultures and induced with 1 mM IPTG at an Abs_600_ ~0.5 and grown at 30 °C for an additional 30 min. Following expression, two buffer C (20 mMTris–HCl (pH 7.5), 50 mM NH4Cl, 25 mM MgCl2) ice cubes were added to each culture flask, rapidly shaken for 1 min on ice and incubated on ice for an additional 30 min. Next, cells were pelleted by centrifugation for 30 min at 4 °C and 10,000×g and resuspended in 600 μL of cold buffer C. Cells were lysed by three cycles of freeze–thawing in liquid nitrogen followed by the addition of three 30-μL aliquots of lysozyme (Novagen), where the stock lysozyme solution was diluted 50-fold in cold buffer C and each lysozyme addition was followed by a 20 min incubation at 4 °C, and finally three additional freeze–thawing cycles. Ribosomes were isolated from these soluble fractions according to previously published procedures ^37,38^. Specifically, three 12-μl aliquots of RQ1DNase (Promega) were added to reduce the viscosity of the lysates and samples were rotated for 15 min at 4 °C after each dose of the enzyme. Samples were spun in a microcentrifuge for 20 min at 10,000×g at 4 °C to pellet the debris. To isolate ribosomes, the supernatant was collected and loaded onto a cold cushion made up of equal volumes of buffer C supplemented with a 5% (w/v) sucrose phase and buffer B (20 mM Tris–HCl (pH 7.5), 500 mM NH4Cl, 25 mM MgCl2) supplemented with a 37% sucrose phase. Ribosomes were isolated by ultracentrifugation for 35 h at 120,000×g and 4 °C using a Beckman LS 8 ultracentrifuge with an SW28 rotor. The crude ribosome pellet was resuspended in 200 μL of buffer C and ultracentrifuged in a 10% to 40% (w/v) sucrose gradient in buffer A (20 nM Tris–HCl (pH 7.5), 100 mM NH4Cl, 25 mM MgCl2) for 17 h at 71,000×g and 4 °C in a SW41 rotor. Gradient fractionation was performed manually by pipetting 250 μL at a time from the top part of the gradient. All collected samples were stored at 4 °C for further analysis. A similar procedure was followed for isolation of ribosomes from cell-free extracts.

### Protein expression and purification

Purification of *Cj*PglB was performed according to a previously described protocol ^31^. Briefly, a single colony of *E. coli* CLM24 carrying plasmid pSN18 ^60^ was grown overnight at 37 °C in 50 mL of LB (10 g/L tryptone, 5 g/L yeast extract, 5 g/L NaCl, pH 7.2) supplemented with ampicillin (Amp) and 0.2% (w/v) D-glucose. Overnight cells were subcultured into 1 L of fresh terrific broth (TB; 12 g/L tryptone, 24 g/L yeast extract, 0.4% (v/v) glycerol, 10% (v/v) 0.17 M KH2PO4/0.72 M K_2_HPO_4_ phosphate buffer), supplemented with Amp and grown until Abs600 reached a value of ~0.7. The incubation temperature was adjusted to 16 °C, after which protein expression was induced by the addition of L-arabinose to a final concentration of 0.02% (w/v). Protein expression was allowed to proceed for 20 h at 16 °C. Cells were harvested by centrifugation and then disrupted using a homogenizer (Avestin C5 EmulsiFlex). The lysate was centrifuged to remove cell debris and the supernatant was ultracentrifuged (100,000×g) for 2 h at 4 °C. The resulting pellet containing the membrane fraction was fully resuspended with a Potter-Elvehjem tissue homogenizer in buffer containing 50 mM HEPES, 250 mM NaCl, 10% (v/v) glycerol, and 1% (w/v) n-dodecyl-β-D-maltoside (DDM) at pH 7.5. The suspension was incubated at room temperature for 1 h to facilitate detergent solubilization of *Cj*PglB from native *E. coli* lipids, which were removed by subsequent ultracentrifugation (100,000×g) for 1 h at 4 °C. The supernatant containing DDM-solubilized *Cj*PglB was purified using Ni-NTA resin (ThermoFisher) according to manufacturer’s specification with the exception that all buffers were supplemented with 1% (w/v) DDM. The elution fraction from Ni-NTA purification was then subjected to size exclusion chromatography (SEC) using an ÄKTA Explorer FPLC system (GE Healthcare) with Superdex 200 10/300 GL column. Purified protein was stored at a final concentration of 1-2 mg/mL in OST storage buffer (50 mM HEPES, 100 mM NaCl, 5% (v/v) glycerol, 0.01% (w/v) DDM, pH 7.5) at 4 °C. Glycerol concentration in the sample was adjusted to 20% (v/v) for long-term storage at −80 °C.

To produce ColE7 for ELISA and affinity enrichment experiments, an overnight culture BL21(DE3) cells carrying plasmid pET28a-ColE7^H569A^ was used to inoculate 1 L of LB supplemented with 50 μg/mL kanamycin. Cells were grown at 37 °C until mid-log phase and then were induced with 0.1 mM IPTG for 16 h at 16 °C before being harvested. Following centrifugation at 10,000×g, pellets were resuspended in PBS buffer supplemented with 10-mM imidazole and lysed at 16,000–18,000 psi using an Emulsiflex-C5 homogenizer (Avestin). The lysate was clarified by centrifugation at 15,000×g for 30 min at 4 °C and the collected soluble fraction was mixed with Ni-NTA resin for 2 h at °C. The mixture was then applied to a gravity flow column and washed with 5 column volumes of PBS containing 20 mM imidazole. Proteins were eluted in 4 column volumes of PBS with 250-mM imidazole. The eluted protein was desalted and concentrated to 5 mg/mL in PBS buffer using Ultracentrifugal Filters with 10-kDa molecular weight cut-off (Amicon) and stored at 4 °C.

### Extraction of LLOs

The protocol for organic solvent extraction of LLOs from *E. coli* membranes was adapted from a previously described protocol ^31^. Briefly, a single colony of strain CLM24 carrying plasmid pMW07-pglΔB for expression of the *C. jejuni N*-glycan on undecaprenylphophate was grown overnight in LB media. Overnight cells were subcultured into 1.0 L of TB supplemented with chloramphenicol and grown until the Abs_600_ reached ~0.7. The incubation temperature was adjusted to 30 °C and protein expression was induced with L-arabinose at a final concentration of 0.2% (w/v). After 16 h, cells were harvested by centrifugation and cell pellets were lyophilized to complete dryness at −70 °C using methanol. For extraction of *Cj*LLOs, the lyophilisates were resuspended in 12 mL of 2:1 (v/v) CHCl_3_:CH_3_OH and sonicated at a 1:1 duty cycle, 5 s cycles, for 100 s. Samples were then centrifuged for 10 min at 3,000×g and the supernatant was decanted, retaining the pellet. The resuspension, sonication, centrifugation, and decanting were then repeated with that pellet. The pellet was then resuspended by vortexing in 20 mL H_2_O, sonicated for 4–5 min until homogenous, centrifuged, and decanted once more. Pellets were then resuspended in 10:20:3 (v/v/v) CHCl_3_:CH_3_OH:H_2_O solution, sonicated until homogenous, and incubated at room temperature for 15 min to facilitate extraction of LLOs. 8 mL of methanol were then added, and samples were centrifuged as before, saving the supernatant while discarding the pellet. The supernatant then had 10 mL of 4:1 (v/v) of chloroform/H2O added and was then centrifuged (4,000×g) for 15 min, after which the organic layer (bottom layer, including flakes) was collected and dried with a vacuum concentrator followed by lyophilization. All lyophilisates containing active *Cj*LLOs were resuspended in cell-free glycosylation buffer (10 mM HEPES, pH 7.5, 10 mM MnCl2, and 0.1% (w/v) DDM) and stored at 4 °C.

### Preparation of crude S12 extracts

CLM24 source strains were grown in 2xYTPG (10 g/L yeast extract, 16 g/L tryptone, 5 g/L NaCl, 7 g/L K_2_HPO_4_, 3 g/L KH_2_PO_4_, 18 g/L glucose, pH 7.2). To generate *Cj*PglB-enriched extract, CLM24 carrying plasmid pSF–*Cj*PglB^61^ was used as the source strain. To generate one-pot extracts containing both *Cj*PglB and *Cj*LLOs, CLM24 carrying pMW07–pglΔB and pSF–*Cj*OST was used as the source strain. After inoculation, the expression of *Cj*PglB was induced at an Abs600 of 0.8 with L-arabinose to a final concentration of 0.02% (w/v). After induction, protein expression was allowed to proceed at 30 °C to a density of Abs_600_ ~3, at which point cells were harvested by centrifugation (5,000×g) at 4 °C for 15 min. All subsequent steps were carried out at 4 °C unless otherwise stated. Cells were harvested and washed twice using S12 buffer (10 mM tris acetate, 14 mM magnesium acetate, 60 mM potassium acetate, pH 8.2). The pellet was then resuspended in 1 mL per 1 g cells of S12 buffer. The resulting suspension was passed once through a EmulsiFlex-B15 high-pressure homogenizer (Avestin) at 20,000-25,000 psi to lyse cells, as we have described previously ^62^. The extract was then centrifuged at 12,000×g for 10 min to remove cell debris and the supernatant was collected and incubated at 37 °C for 60 min shaking at 250 rpm for the run-off reaction. Following centrifugation at 10,000×g for 10 min at 4 °C, the supernatant was collected, flash-frozen in liquid nitrogen, and stored at –80 °C.

### *In vitro* reconstituted glycosylation and cell-free glycoprotein synthesis

For *in vitro* reconstituted glycosylation of *E. coli-cell* derived RNC complexes displaying Im7^N58^ or ^DQNAT^scFv13–R4, reactions were carried out in a 50-μL volume containing 3 μg of ribosome-stalled acceptor protein, 2 μg of purified *Cj*PglB, and 5 μg extracted LLOs in *in vitro* glycosylation buffer (10 mM HEPES, pH 7.5, 10 mM MnCl2, and 0.1% (w/v) DDM). The reaction mixture was incubated at 30 °C for 16 h. For CFGpS, reactions were carried out in 1-mL reaction volumes in a 15-mL conical tube using a modified PANOx-SP system ^62^. The reaction mixture contained the following components: 0.85 mM each of GTP, UTP, and CTP, 1.2 mM ATP, 34.0 μg/ml folinic acid, 170.0 μg/ml of *E. coli* tRNA mixture, 130 mM potassium glutamate, 10 mM ammonium glutamate, 12 mM magnesium glutamate, 2 mM each of 20 amino acids, 0.4 mM nicotinamide adenine dinucleotide (NAD), 0.27 mM coenzyme-A (CoA), 1.5 mM spermidine, 1 mM putrescine, 4 mM sodium oxalate, 33 mM phosphoenolpyruvate (PEP), 57 mM HEPES, 6.67 μg/ml plasmid, and 27% (v/v) of cell lysate ^63^. Protein synthesis was carried out for 30 min at 30 °C, after which protein glycosylation was initiated by the addition of MnCl_2_ and DDM at a final concentration of 10 mM and 0.1% (w/v), respectively, and allowed to proceed at 30 °C for 16 h. Isolation of ribosomes from *in vitro* reconstituted glycosylation and CFGpS reactions was performed as described above.

### Western blot analysis

To concentrate ribosome fractions for Western blot analysis, volumes of the collected ribosomal fractions were mixed with cold 20% (v/v) trichloroacetic acid (TCA) in a 1:2 volume ratio and allowed to precipitate for 30 min on ice. After ultracentrifugation for 20 min at 210,000 and 4 °C, the pellet was dried of all remaining TCA and directly resuspended in SDS–PAGE loading buffer. Ribosome fractions were resolved by SDS-polyacrylamide gel electrophoresis on 10% Mini-PROTEAN TGX™ Precast Protein Gels (Bio-Rad). Prior to loading, fractions were normalized by rRNA content as measured by A260 (see **Supplementary Figs. 1** and **2**). For visualizing the separated protein samples, gels were stained with Coomassie G-250 stain (Bio-Rad) following the manufacturer’s protocol. For Western blot analysis, the separated protein samples were then transferred to nitrocellulose membranes using a semi-dry apparatus. Following transfer, the membranes were blocked with 5% milk (w/v) in TBST (1x TBS, 0.1% Tween 20) and subsequently probed for 1 h with one of the following: horseradish peroxidase (HRP)-conjugated anti-DDDDK antibody (Abcam, cat # ab49763) that recognized the FLAG epitope tag; the *C. jejuni* heptasaccharide glycan-specific antiserum hR6 (kindly provided by Markus Aebi); or the mouse mAb FB11 (ThermoFisher; cat # MA1-7388) that specifically recognizes *F. tularensis* LPS. Goat antirabbit IgG (HRP) (Abcam, cat # ab205718) was used as the secondary antibody to detect hR6 antiserum while HRP-conjugated anti-mouse IgG (Abcam, cat # ab97023) was used as the secondary antibody to detect FB11. After washing five to six times with TBST for 5 min, the membranes were visualized using a ChemiDoc MP Imaging System (Bio-Rad).

### ELISA

Binding activity for ribosome-tethered Im7^N58^ and scFv–HER2^DQNAT^ was determined by standard ELISA. Briefly, treated Costar 96-well ELISA plates (Corning) were coated overnight at 4 °C with 50 μL of 5-μg/mL in-house prepared ColE7 in PBS for ribosome-tethered Im7^N58^ or the same amount of commercial extracellular domain (residues 1 to 652) of human HER2 (HER2-ED; Sino Biological, 10004-HCCH) for ribosome-tethered scFv–HER2^DQNAT^ in PBS. After blocking with 200 μL of blocking solution (1% (w/v) non-fat milk, 5 mM MgCl_2_, 2.5 mg/ml of heparin, 0.05 mg/ml *E. coli* tRNA in PBS), for 2 h at room temperature, the plates were washed four times with washing solution (0.1% (v/v) Tween 20, 5 mM MgCl_2_ in PBS). Isolated 70 S ribosome samples displaying Im7^N58^ and scFv–HER2^DQNAT^ were mixed gently with equal volume of cold blocking solution, and 100 μl of this mixture was added to each well. The plate was incubated for 1 h at 4 °C and then washed five to six times with 200 μl of cold washing solution at 4 °C to remove any unbound complexes. After washing, 50 μL of the HRP-conjugated anti-DDDK antibody (see above) in 1% PBST was added to each well for 1 h. Plates were washed three times and then developed using 50 μL 1-Step Ultra TMB-ELISA substrate solution (ThermoFisher). A similar ELISA protocol was followed for detecting RNC complexes displaying glycosylated ^DQNAT^pd except that the plates were coated with 50 μL of 5-μg/mL mouse mAb FB11 in PBS and were subsequently detected with HRP-conjugated anti-mouse IgG.

### Affinity selection of ribosome complexes, mRNA isolation, and RT-PCR

Affinity selection was performed using ELISA plates that were prepared and treated with 70S ribosome samples identically as described above. However, instead of detection of bound complexes with antibodies, mRNA was dissociated from bound ribosome complexes by adding 100 μl of cold eluting solution (20 mM EDTA, 20 units/ml RNasin in PBS) to each well and shaking gently for 30 min at 4 °C. Samples were collected into cold microcentrifuge tubes after scraping the plate surface with a tip to ensure complete sample removal. mRNA was purified using the RNeasy Purification Kit (Qiagen). RT-PCR on the recovered mRNA was performed using the SensiScript RT Kit (Qiagen) with reverse primers that bound the 3’ end of each POI sequence. PCR amplification was performed in a second step with the same reverse primer and a forward primer specific for the 5’ end of each POI sequence. The PCR products were either submitted directly for DNA sequencing or instead ligated into plasmid pET28a, which was then submitted for DNA sequencing to confirm the identity of the glycosylated POI stalled on ribosomes.

## ACKNOWLEDGMENTS

We thank Markus Aebi for providing strain CLM24 and hR6 serum used in this work. We also thank Peter Schweitzer and the BRC Genomics Facility (RRID:SCR_021727) at the Cornell Institute of Biotechnology for sequencing experiments. This work was supported by the Defense Threat Reduction Agency (HDTRA1-15-10052 and HDTRA1-20-10004 to M.P.D. and M.C.J), the Bill and Melinda Gates Foundation (OPP1217652), and the NSF (CBET-1159581 and CBET-1264701 to M.P.D. and CBET-1936823 to M.P.D and M.C.J.). E.J.B. was supported by a training grant from the NIH/NIGMS Chemical Biology Interface Training Grant (T32GM138826) and an NSF Graduate Research Fellowship (DGE-2139899). J.M.H. and K.F.W were supported by NDSEG Fellowships.

## AUTHOR CONTRIBUTIONS

S.S.C. and E.J.B. designed research, performed research, analyzed data, and wrote the paper. J.M.H. and K.F.W. designed research, performed research, and analyzed data. M.C.J. and M.P.D. designed and directed research, analyzed data, and wrote the paper.

