## Supplementary Figures 1 and 2 for "Ribosome display of *N*-linked glycoproteins in cell-free extracts"

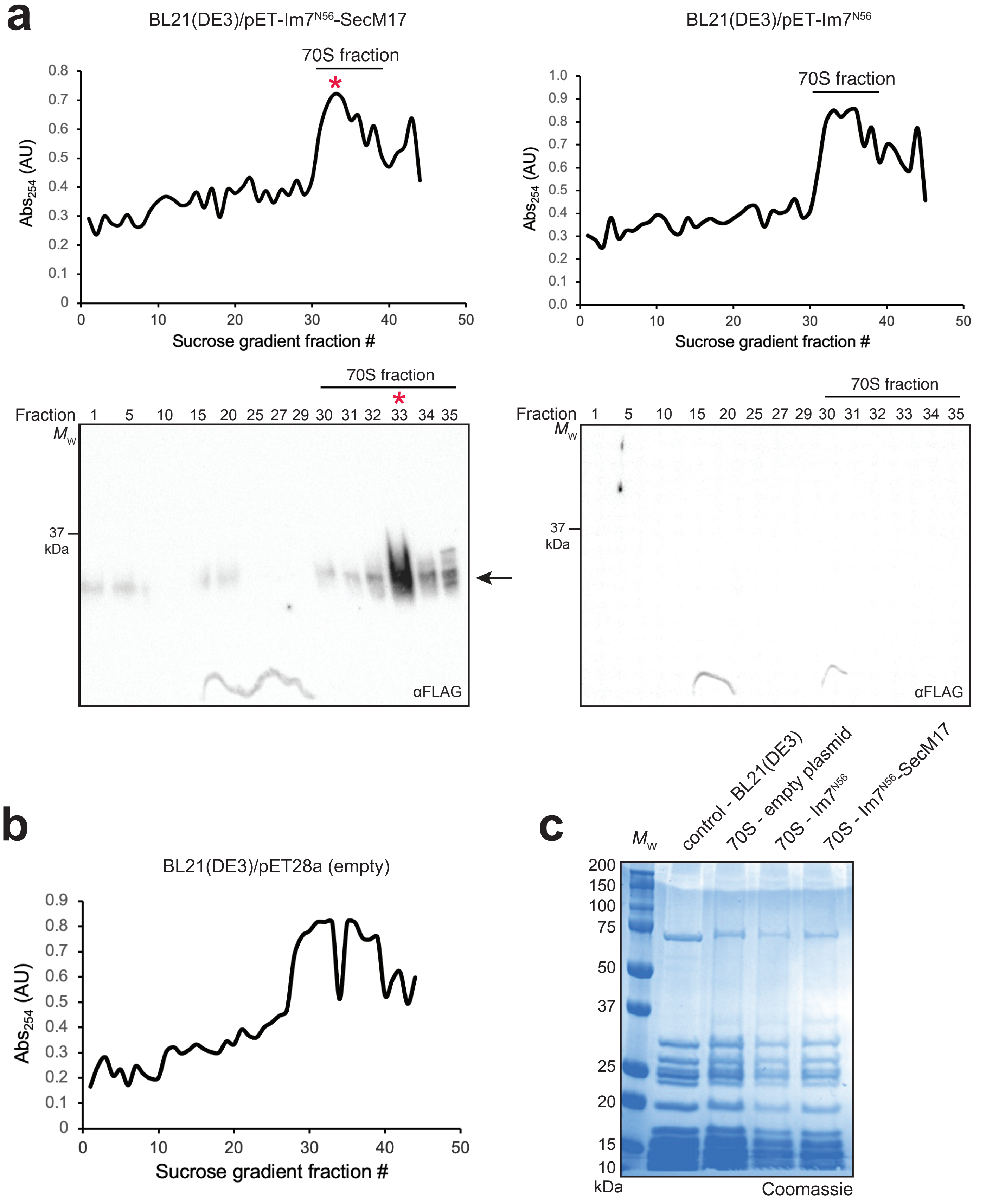


**Supplementary Figure 1. Purification and characterization of 70S ribosomes displaying Im7$^{N56}$-SecM17.** (a, top) Absorbance trace (254 nm) of sucrose fractions derived from BL21(DE3) cells expressing Im7$^{N56}$-SecM17. Red asterisk indicates the fraction selected for further downstream analysis and used in hybrid glycoRNC reactions. (a, bottom) Western blot analysis of selected 70S ribosome sucrose fractions from (a, top) as indicated. Blots were probed with α-FLAG-HRP antibody specific for epitope tag on Im7$^{N56}$-SecM17. Black arrow indicates expected molecular weight of Im7$^{N56}$-SecM17. (b) Absorbance trace (254 nm) of sucrose fractions derived from BL21(DE3) cells carrying empty pET28a plasmid. (c) Coomassie-stained SDS-PAGE gel of sucrose fractions enriched with 70S ribosomes from BL21(DE3) cells without a plasmid, with empty pET28a plasmid, expressing Im7$^{N56}$, and expressing Im7$^{N56}$-SecM17. Molecular weight ($M_W$) markers are indicated on left of Western blots and SDS-PAGE gel. All results are representative of biological replicates ($n = 2$).

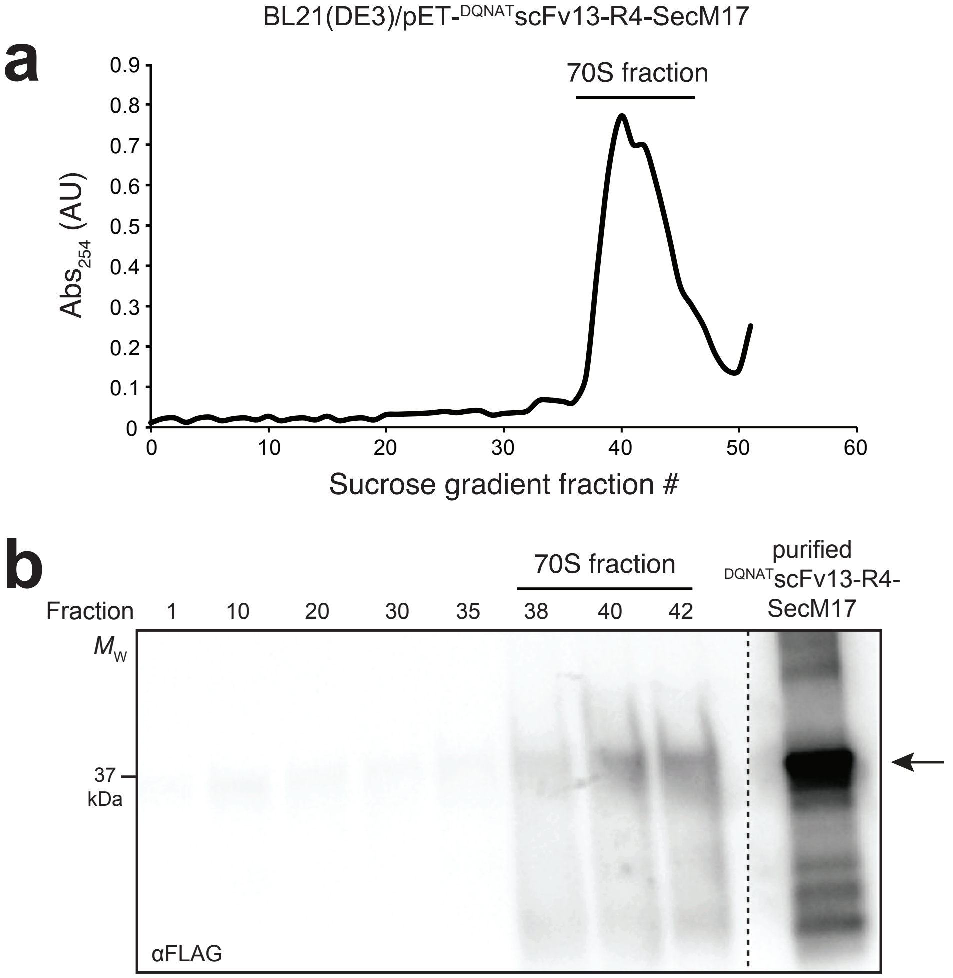


**Supplementary Figure 2. Purification and characterization of 70S ribosomes displaying $^{DQNAT}$scFv13-R4-SecM17.** (a) Absorbance trace (254 nm) of sucrose fractions derived from BL21(DE3) cells expressing $^{DQNAT}$scFv13-R4-SecM17. Results are representative of biological replicates ($n$ = 2). (b) Western blot analysis of selected 70S ribosome sucrose fractions from (a) as indicated. Blots were probed with α-FLAG-HRP antibody specific for epitope tag on $^{DQNAT}$scFv13-R4-SecM17. Black arrow indicates the expected molecular weight of $^{DQNAT}$scFv13-R4-SecM17, as demonstrated by the purified control at right. Molecular weight ($M_W$) markers are indicated on left. Western blot is representative of biological replicates ($n$ = 2).
